# Eco-physiological and transcriptomic plasticity of *Dianthus inoxianus* in response to drought

**DOI:** 10.64898/2026.04.08.702570

**Authors:** Alba Rodríguez-Parra, Francisco Balao

## Abstract

Phenotypic plasticity is a key mechanism by which plants adjust their traits to environmental changes. These phenotypic adjustments are driven by plastic changes in gene expression regulated by gene regulatory networks. Drought, a major selective force in Mediterranean ecosystems, provides a powerful context to examine how genomic plasticity translates into phenotypic responses. Here, we used *Dianthus inoxianus*, a drought-tolerant Mediterranean carnation, in order to characterize the phenotypic and transcriptomic plasticity in response to drought stress combining ecophysiological measurements with RNA-seq, gene co-expression and gene regulatory network analyses. Most of the phenotypic traits exhibited low plasticity in response to drought, except water and osmotic potential. At transcriptome level, we identified 57 plastic genes, suggesting that drought tolerance in *D. inoxianus* relies predominantly on constitutive gene expression. These plastic genes were enriched in processes typically related to drought response, such as cell wall components and abscisic acid (ABA) signaling. Some plastic genes belonged to drought-responsive modules, while others were hubs in different modules acting as inter-modular connectors. Furthermore, the regulatory network revealed that these plastic genes were strongly regulated by multiple stress-responsive transcription factors, and that drought-associated modules were regulated through both ABA-dependent and ABA-independent pathways. In addition, we identified contrasting patterns of canalization and decanalization, with immune and post-transcriptional regulation remaining canalized under drought, whereas photosynthesis and amino acid metabolism became decanalized, potentially releasing cryptic genetic variation. Overall, our results emphasise that drought tolerance in *D. inoxianus* emerges from a strategy combining preadaptation with targeted plasticity in key molecular pathways.

## 1. Introduction

Phenotypic plasticity, the ability of an organism to express different phenotypes in response to changes in the environment, has been recognized as one of the main mechanisms by which plants will respond to rapid environmental changes (Bradshaw, 1965; Valladares et al., 2007). The environment can influence phenotypes in numerous diverse and complex ways, causing changes in important traits ranging from morphology, physiology and anatomy to life history traits (Caswell, 1983; Gratani et al., 2006; Navas & Garnier, 2002). This phenotypic flexibility represents one of the main short-term responses of plant populations to global change scenarios (Gibert et al., 2019; Matesanz et al., 2010). By adjusting key functional traits in response to these novel pressures, phenotypic plasticity can enhance plant survival and performance, acting as an important buffer against environmental unpredictability (Nicotra et al., 2010).

Among the major environmental stressors shaping plant performance, drought stands out as one of the most critical according to current and projected climate change scenarios (Cook et al., 2018). This is particularly evident in the Mediterranean region which is characterized by high temperatures and prolonged summer droughts (Deitch et al., 2017; Schär et al., 2004), affecting nearly all aspects of plant physiology and metabolism. Consequently, plants coordinate their adaptive responses through multiple mechanisms (Farooq et al., 2012), including physiological adjustments. In this context, plasticity in physiological traits enables plants to modulate their metabolism and acclimate to adverse conditions (Gratani, 2014). Indeed, several studies have documented the plasticity of leaf physiological traits, such as water potential (Johnson et al., 2018), osmotic potential (Cardoso et al., 2018) and stomatal density, as well as in functional traits including specific leaf area (SLA; De Kort et al., 2020) and leaf dry matter content (LDMC; Pérez-Ramos et al., 2019) in response to drought. Additionally, drought response is complex mechanism from the genomic perspective. This complexity arises because drought resistance entails a suite of physiological, morphological, and biochemical adaptations, each controlled by different sets of genes (Blum, 2011). Drought stress causes cellular damage and induces osmotic and oxidative stress. This triggers a signaling cascade that involves transcription factors which activate stress response mechanisms. From an adaptive point of view, these stress response mechanisms can be divided into three broad groups: 1) osmotic adjustment, 2) damage control and repair, and 3) growth regulation (Roldán-Arjona & Ariza, 2009; Serraj & Sinclair, 2002; Wang et al., 2003). Moreover, the molecular control mechanism of stress is mediated by: (i) Signaling and transcriptional control regulators, including PP2Cs, MAPK cascades, SOS/CBL–CIPK kinases, phospholipases, and major drought responsive transcription factor families such as HSF, CBF/DREB, and ABF/AREB, coordinate early stress perception and integrate ABA dependent and ABA independent pathways to reprogram gene expression, thereby inducing protective functions related to osmotic adjustment, ROS detoxification, and cellular protection (Manna et al., 2021; hereafter dTFs); (ii) Genes involved in the exchange and transport of water and ions, like aquaporins (Li et al., 2025) and ion transporters, as well as genes involved in the recovery of homeostasis (stomatal control or photosynthesis); (iii) Genes involved in the protection of membranes and proteins, such as heat shock proteins (Hsps), chaperones, and LEA; and (iv) Genes involved in cell wall growth and remodeling (Moore et al., 2008), which regulate cell expansion and mechanical properties through pathways controlling wall loosening and polysaccharide deposition, thereby constraining growth under water limitation and contributing to stress tolerance; and (v) Growth regulation genes, such as those involved in the biosynthesis of abscisic acid (Xiong & Zhu, 2003). However, these mechanisms have been dilucidated in model species, and it remains uncertain how they translate to non-model organisms, which often exhibit different ecological and genetic complexities.

In this regard, understanding the role of plasticity in evolution starts at the level of gene expression (Sultan, 2021) which is a crucial intermediary in the evolutionary interplay between organisms and their environments (Rivera et al., 2021). Gene expression plasticity (GEP), which concerns the capacity of genes to change their expression levels under diverse conditions, is critical for phenotypic plasticity, adaptation and evolvability (Rivera et al., 2021). GEP allows for immediate and reversible adjustments in phenotype without altering its underlying DNA sequence. For instance, in the presence of stressors such as temperature fluctuations, water scarcity, or pathogens, cells can rapidly adjust the expression levels of certain genes (De Nadal et al., 2011). These adjustments can lead to changes in the production of proteins that are critical for the immediate response and adaptation to these stressors. However, it remains an open question whether insights from the plasticity of gene expression levels can be applied to more complex phenotypic traits (Chevin et al., 2022). GEP is controlled by complex gene regulatory networks (GRNs) that integrate signals from both internal and external environments to modulate gene expression. The flexibility and responsiveness of these networks are key to fitness and evolutionary success. GRNs are particularly important in understanding phenotypic plasticity as they control the timing, localization and level of gene expression, and their flexibility allows organisms to adjust their phenotype to their environment. For example, in response to temperature changes or nutrient availability, GRNs can activate or repress specific genes, leading to changes in the phenotype (MacNeil & Walhout, 2011).

Although phenotypic plasticity is generally understood as changes in the mean value of a trait across contrasting environments, plasticity can also occur in trait variance due to underlying genotypic diversity. In this context, “canalization” refers to the robustness of phenotypes to perturbations, meaning that gene expression would remain relatively stable despite external (environmental) or internal (genetic) disturbances (Waddington, 1942). Therefore, under strong canalization, variance in plasticity is low. In contrast, “decanalization” arises when genotypes respond differently to environmental factors, increasing variance in plasticity and revealing previously hidden genetic differences (Whitman & Agrawal, 2009). Consequently, decanalization can unmask phenotypic variation that may be subject to selection, thereby enhancing the evolvability of the trait within a population (Flatt, 2005). In this way, environmentally induced responses that are initially reversible could, under persistent selection, become fixed adaptive trait (Crispo, 2007; Levis & Pfennig, 2016).

In this study, we assess the phenotypic and transcriptomic plasticity of *Dianthus inoxianus* Gallego (Caryophyllaceae) in response to drought stress. This species is a polyploid taxon belonging to the *Dianthus broteri* Boiss. & Reuter complex (Balao et al., 2009, 2010). *D. inoxianus* is a perennial herbaceous plant endemic to Doñana National Park, located in the lower Guadalquivir River valley (Balao et al., 2009). This wild carnation typically occupies narrow ecological niches on nutrient-poor and dry sandy soils, where it is exposed to pronounced seasonal drought (Balao et al., 2009; López-Jurado et al., 2019). Previous studies have shown that *D. inoxianus* exhibits eco-physiological adaptations that confer high drought tolerance, including a remarkable recovery capacity (López-Jurado et al., 2016, 2020) and highly competitive capacity (Rodríguez-Parra et al., 2025) following water shortage. Notably, *D. inoxianus* has been reported to follow an acquisitive strategy, restricting its growing season to brief windows of opportunity when water and nutrients are sufficiently available (López-Jurado et al., 2022). Moreover, transcriptomic studies have revealed a rapid response to thermal shock through changes in the expression of several stress-responsive genes (e.g., *SWEET1*, *PP2C16*, *ATHB7*) involved in high-temperature responses (López-Jurado et al., 2024), although these genes have also been described as responsive to water deficit (Chen et al., 2022; Pruthvi et al., 2014; Zhou et al., 2017). Altogether, these findings suggest that in addition to its inherent eco-physiological adaptations, *D. inoxianus* may have evolved an enhanced ability to cope with drought stress through phenotypic or genetic plastic responses. Alternatively, this drought tolerance may be driven by the constitutive expression of stress-responsive genes rather than by environmentally induced plasticity (Steele et al., 2025), highlighting the need to disentangle whether drought tolerance in *D. inoxianus* is fundamentally plastic, constitutive, or the result of both mechanisms.

In this context, the present study aims to provide a characterization of the phenotypic and transcriptomic plasticity of *Dianthus inoxianus* under drought stress, elucidating the functional and regulatory mechanisms that enable this species to cope with water scarcity. To address this, we (1) quantify phenotypic plasticity in eco-physiological traits, (2) identify the transcriptome response to drought and key plastic genes involved, and (3) infer gene regulation networks associated with drought stress.

## 2. Material & Methods

### 2.1 Experimental set-up and plant sampling

We used two clonal plants derived from apical cuttings of 10 different genotypes of *Dianthus inoxianus* (n=20) from three different natural populations. To promote root development, the cuttings were treated with indole-3-butyric acid (IBA, 500 mg/L) and 1-naphthaleneacetic acid (NAA, 500 mg/L) solutions and subsequently maintained in an aeroponic system (see López-Jurado 2019 for further details). After the rooting of cuttings, plants were transplanted into 2.5 L pots filled with a commercial organic substrate (Gramoflor GmbH und Co. KG.) and perlite mixture (3:1) inside the General Greenhouse Services of the University of Seville, maintaining controlled temperature conditions at 21/25 °C, 40-60% relative humidity and a maximum photosynthetic photon flux density (PPFD) incident on leaves of 1200 μmol m^-2^ s^-1^ during 12 months.

At the beginning of the experiment, all plants were watered to field capacity and then were transferred to a FITOCLIMA 18000 EH culture chamber (Aralab, Portugal) arranged in a randomized plot with a 14:10h photoperiod, controlled temperature of 25°C/14°C, and a relative humidity of 60%. Then they were randomly divided into two distinct water regimes (10 clonal plants in each condition). One group was consistently kept under well-watered conditions (WW) by irrigating daily to field capacity, while the other group was subjected to water stress (WS) by suspending irrigation until severe water stress was reached. During the experiment, we monitored the soil water content (SWC) daily, to track its progressive decline. Severe water stress was reached on day 18, defined as the point when stomatal conductance to CO₂ (gₛ) dropped below 50 mmol H₂O m⁻² s⁻¹ (Medrano et al., 2002), a threshold indicating that photosynthetic activity becomes primarily limited by metabolic constraints.

### 2.2 Phenotypic plasticity of functional traits in response to drought stress

Throughout the experiment, a set of ecophysiological traits were measured at the beginning of the experiment (1^st^ day) and at the severe water stress day (18^th^ day). To evaluate leaf water status, we quantified the total leaf water potential (Ψw), osmotic potential (Ψπ), and relative water content (RWC) in fully developed random leaves in each water regimes (n = 4 for Ψw and Ψπ; n=6 for RWC). These measurements were conducted twice at day: at pre-dawn (Ψw_pd_, Ψπ_pd_, and RWC_pd_) and at midday (Ψw_md_, Ψπ_md_, and RWC_md_). Ψw was determined using a pressure chamber (Model 1515 D - Pressure Chamber Instruments, PMS), while Ψπ was measured via the psychrometric technique using a vapor pressure osmometer (5600 Vapro, Wescor, Logan, USA). RWC was calculated as follows:

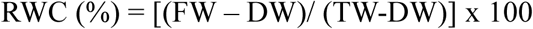

where FW represents the fresh weight of the leaf, DW was the dry weight of the leaf after drying at 60 °C in the oven to constant weight, and TW was the saturated weight of the leaf after 24 h of rehydration in distilled water at 4 °C in darkness to minimize respiration losses.

To assess the effects of severe drought on photochemical efficiency, we investigated the maximum quantum efficiency of PSII photochemistry (*F*_v_/*F*_m_) using a pulse-modulated fluorometer (FMS-2, Hansatech Instrument Ltd., UK) with dark-adapted leaves (n = 10 per water regime) under saturating light intensity at midday (1600 μmol photon m⁻² s⁻¹).

To evaluate the phenotypic plasticity of functional traits, we used the relative distance plasticity index (RDPI;Valladares et al., 2006). Plasticity was calculated on the day of severe drought as the trait distance (in absolute value) of the same genotypes between the two water regimes (WW-WS) divided by the average of the two genotypic trait values. Then, all individual RDPI values for each phenotypic trait were averaged to obtain a single mean RDPI value per trait. All these statistical analyses were carried out in R ver. 4.3.3 (R Core Team, 2024).

### 2.3 RNA Extraction, cDNA Library Preparation, and Illumina Sequencing

Fully developed leaves were collected from both WW and WS plants (n = 6 per regime) on the day of severe drought stress and were immediately frozen in liquid nitrogen and stored at −80°C until processing. RNA was then extracted from the leaves using the Direct-zol RNA Miniprep commercial kit (Zymo Research, Irvine, CA, USA) in accordance with manufacturer’s instructions. Subsequently, RNA sequencing libraries were constructed from the isolated RNA using the TruSeq Stranded mRNA LT Sample Prep Kit. Libraries were sequenced using Illumina NovaSeq 6000 as 150 bp pair-end. Raw reads were initially processed with Fastp v0.23.3 to remove adapters and trim low-quality regions. Subsequent filtering of rRNA contamination was filtered using SortMeRNA v4.3.2. The resulting clean reads were deposited in the NCBI Short Reads Archives (SRA; BioProject ID PRJNA1248684).

### 2.4 Identification of plastic genes associated with drought

The quality-trimmed RNA-seq reads were aligned to the 2x *D. broteri* reference genome (Picazo-Aragonés *et al*., 2025, manuscript under review) with STAR v. 2.7.10b (Dobin et al., 2013) using the gene models annotated. The following STAR parameters were applied: “-outFilterScoreMinOverLread” 0.1, “-outFilterMatchNMinoverLread” 0.1 and “-outFilterMismatchNmax” 5.

To investigate the genes with plasticity under drought response, we modeled gene expressions as negative binomial mixed models using glmmSeq (Lewis et al., 2025), with watering treatment (WW vs WS) as fixed effect and clone identity as a random intercept. Counts were normalized using TMM and filtered to retain genes with CPM ≥ 1 in at least three samples. Gene-wise dispersions were estimated with edgeR (Robinson et al., 2009) and supplied to glmmSeq, and q-values were obtained using the Benjamini–Hochberg procedure. The functional enrichment of the plastic genes was assessed through Gene Ontology (GO) enrichment analysis. The genome annotation was imported into R and analyzed with *topGO* v.2.60.1 (Alexa & Rahnenführer, 2022). Using the classic algorithm with Fisher’s exact test, we retained GO terms with an adjusted p-value (FDR ≤ 0.05). Additionally, the resulting enriched GO terms were summarized and visualized in R.

### 2.5 Canalization of gene expression in response to drought stress

To test whether drought affected the variance of gene expression (canalization and decanalization), we fitted for each gene two alternative negative binomial models using the R package glmmTMB (Brooks et al., 2017) with a negative binomial distribution and quadratic variance parameterization (Hardin & Hilbe 2007). The first model assumed a single dispersion parameter shared across treatments (i.e. WW and WS), whereas the second model allowed dispersion to vary between treatments. Models were fitted gene-by-gene, using a BFGS optimizer, and we compared the constant-vs treatment-specific dispersion models for each gene with a likelihood ratio test (LRT). For genes with LRT p-value < 0.01, we computed the difference in dispersion estimates to classify genes as canalized under drought (lower variance in drought) and decanalized under drought (higher variance in drought). Genes with non-significant LRT were considered canalized across treatments (no detectable change in variance). To visualize changes in expression variance between treatments, we estimated gene-wise biological coefficients of variation (BCV) separately for each treatment using edgeR. Tagwise dispersions were then estimated independently in each treatment, and BCV values were obtained as the square root of the tagwise negative binomial dispersion for each gene. Finally, to identify the biological functions in which the canalized and decanalized genes were involved, we performed a GO enrichment analysis, using the parameters described above.

### 2.6 Gene Coexpression and identification of gene modules associated with drought response

We constructed gene co-expression networks (GCNs) using *BioNERO* (Almeida-Silva & Venancio, 2022). Prior to network inference, expression matrices were filtered using the *percentage-based* approach. We filtered out genes with fewer than one count in at least 50% of samples and applied a variance-stabilizing transformation (Prost-Boxoen et al., 2025), using a more permissive filter than for plasticity tests to retain lowly expressed but variable genes. The GCN was inferred using a signed hybrid adjacency, biweight midcorrelation, and a minimum module size of 50 genes. Approximate scale-free topology was assessed by selecting a soft-thresholding power as the first stable plateau yielding an acceptable scale-free fit (Supplementary Fig. 1). Module detection combined hierarchical clustering of the topological overlap matrix with dynamic tree cutting, as implemented in BioNERO.

We identify drought-responsive modules by correlating module eigengenes with drought treatment. Modules showing significant associations (FDR ≤ 0.05) were considered drought responsive, and hub genes within each of these modules were considered as those showing both high module membership (|kME| > 0.8) and high intramodular connectivity (top 10%). Functional enrichment for each module was assessed using GO terms, as previously described.

### 2.7 Integrated Gene Regulation Network of plasticity in response to drought

A total of 1,616 TFs were identified from the 2x *D. broteri* genome with PlantTFDB 5.0 (Tian et al., 2020). Putative TFs were identified using BLASTp with a cut-off *E*-value of 1 × 10^−5^ and best hit in *Arabidopsis thaliana*. Of these, 1,362 passed the expression filtering and were retained for downstream analyses. GRNs were inferred using the GENIE3 algorithm (Huynh-Thu et al., 2010) implemented in BioNERO, with all annotated transcription factors and an ensemble of 10,000 regression trees. A GRN inference using the same expression filtering applied for the GCN, based on the full dataset (12 samples: 6 WW and 6 WS), allowing the model to capture regulatory structure emerging from variation across both conditions. To obtain robust scale-free subnetwork, we applied a constrained selection based on topology and stability. Subnetworks were built across increasing edge-weight quantiles, and for each we calculated the scale-free fit (R²) and structural stability between consecutive quantiles (Jaccard index). We selected the 94th edge-weight quantile as the optimal threshold (R² = 0.60, Jaccard = 0.92; ∼2.43M edges), prioritizing the stable R² plateau over a sparse local maximum at higher quantiles.

We computed global network centralities on the filtered GRN using out-degree, betweenness and directed closeness of the TFs to identify global regulators occupying high-connectivity and high-mediation positions within the network. Each metric was standardized (z-scores) and summarized in a composite centrality score (mean z across metrics). We then examined the regulatory contribution of all TFs and drought-responsive TFs (Manna et al., 2021; dTFs afterwards; Supplementary Table 3) to (i) the 57 plastic genes and (ii) the 10 GCN modules associated with drought. The filtered GRN was integrated with the GCN by assigning each TF–target interaction to the module of the target gene, allowing us to quantify the regulatory influence of each TF across modules. In addition, we ranked TFs within each drought-associated module using a score combining statistical support and module specificity (−log₁₀(p-adjusted) × fraction of targets in the module) and selected the top 10 TFs per module to identify the strongest module-level regulators independently of the previously defined drought-responsive TFs. For each TF, we calculated the proportion of its targets belonging to each module and tested for over-representation relative to the genome-wide module distribution (Fisher’s exact test; FDR < 0.05). This identified TFs exerting module-focused regulatory activity. Regulatory load at the module level was quantified as the mean incoming edge weight per gene, calculated as the total weight of all TF–target connections associated with a module divided by the number of genes it contains. Because GENIE3 weights represent the importance of each regulatory interaction, this metric provides a size-independent estimate of the aggregated regulatory intensity acting on each module. Finally, TF regulatory profiles were visualized using bipartite graphs linking TFs to the modules they predominantly regulate.

## 3. Results

### 3.1 Phenotypic plasticity in functional traits

In this study, the phenotypic traits showed substantial variability. Ψw_pd_ and Ψw_md_ ranged from –0.24 MPa to –1.53 MPa and from –0.56 MPa to –1.80 MPa, respectively. Ψπ_pd_ and Ψπ_md_ varied between –1.36 MPa and –2.44 MPa, and –1.71 MPa and –2.86 MPa, respectively. RWC_pd_ and RWC_md_ ranged from 55.76 to 85.45%, and from 53.51 to 74.25%, respectively, while Fv/Fm values ranged from 0.383 to 0.741. Some of the measured traits exhibited significant differences between WW and WS conditions such as Ψw_pd_, Ψπ_md_ and RWC_md_. In particular, Ψw_pd_ was the trait most affected by drought (mean ± SD; WW: – 0.548 ± 0.235 MPa vs. WS: –1.34 ± 0.487 MPa; p. value = 0.005 ;Fig. 1b), followed by Ψπ_md_ (WW: –1.92 ± 0.221 MPa vs. WS: –2.28 ± 0.399 MPa; p. value= 0.095; Fig. 1c) and RWC_md_ (WW: 66.6 ± 6.87% vs. WS: 59.8 ± 5.84%; p. value= 0.095; Fig. 1d; see Supplementary Table 1 for details on the remaining traits). Not all traits exhibited the same degree of plasticity, with an overall RDPI of 0.151 ± 0.133 (Fig. 1a). The highest plasticity was detected for Ψw_pd_ (RDPI 0.439 ± 0.222), followed by Ψw_pd_ (0.176 ± 0.105) and Ψπmd (0.136 ± 0.072). In contrast, RWCpd exhibited the lowest plasticity (0.053 ± 0.039).

**Figure 1:**
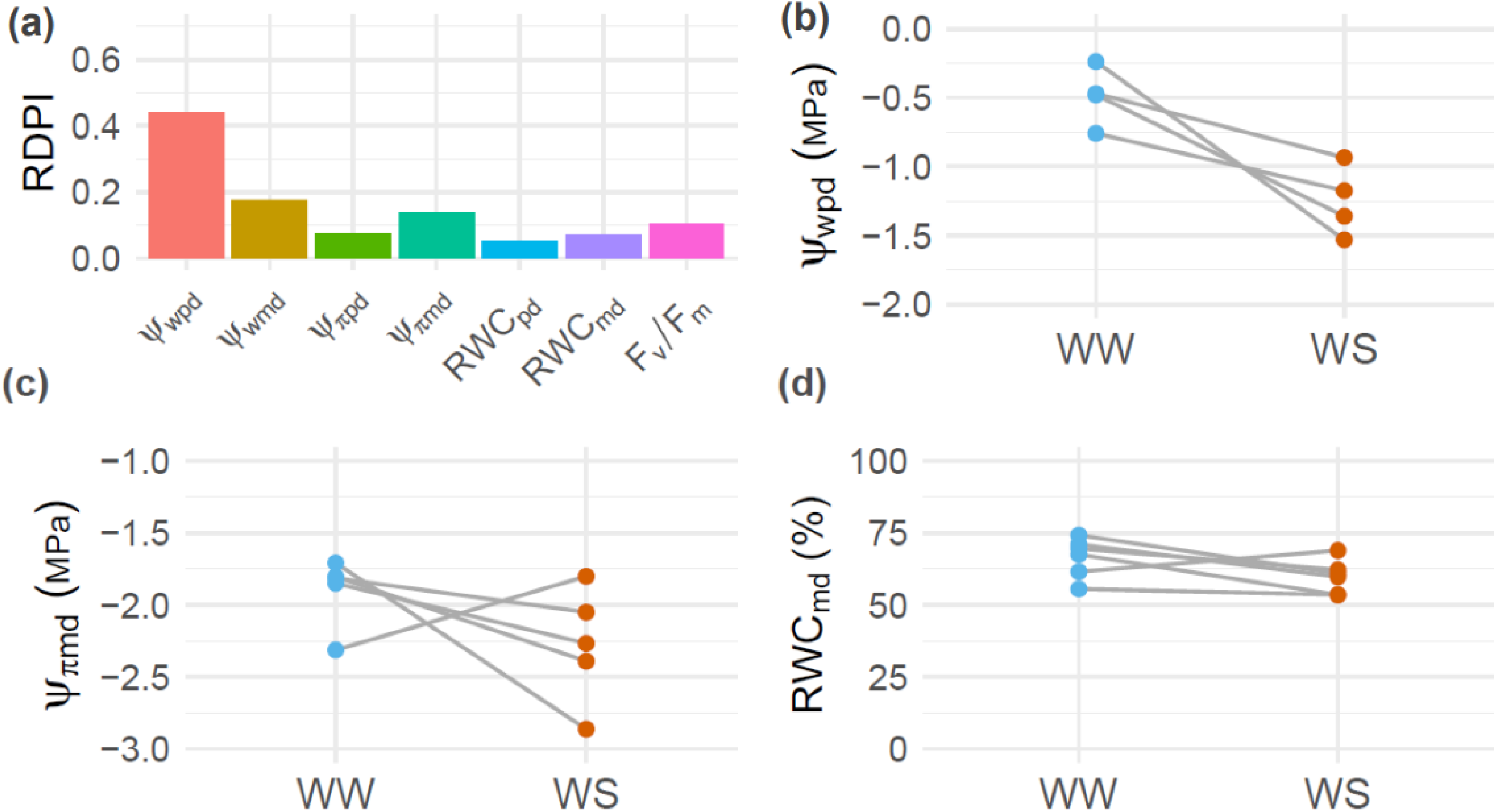
Trait plasticity in response to drought stress. (a) Relative Distance Plasticity Index (RDPI) of physiological traits. (b) Predawn water potential (Ψw_pd_) under well-watered (WW) and water-stressed (WS) conditions. (c) Midday osmotic potential (Ψπ_md_) values. (d) Midday relative water content (RWC_md_) values.

### 3.2 Transcriptomic plasticity genes

The RNA sequencing resulted in an average of 810 ± 28 M reads per sample. Read mapping rate to the reference genome ranged from 99.30% to 99.82%. After normalization of expression and gene filtering, a total of 23,700 genes were expressed. However, the total number of genes expressed within each treatment was similar (WW: 22,160 and WS: 22,145 genes).

The GLMM analyses identified 57 genes exhibiting significant expression plasticity in response to drought (p-adjusted < 0.05). Although not all genes responded in the same direction (Supplementary Table 2), 25 genes showed downregulation under drought stress, such as DIABRO_002784 (Fig. 2a), which is involved in calcium ion transport to the mitochondria (MCU3-like). Conversely, 32 genes displayed upregulation under drought stress, as exemplified by DIABRO_023861 (ATCESA7-like; Fig. 2b) associated with cellulose synthesis, a key component of the cell wall. The functional enrichment analysis of the 57 plastic genes (Fig. 2c) revealed that drought significantly altered functions as cell wall component biogenesis (GO:0030243, GO:0051273) and mitotic cell cycle (GO:0000278). Additionally, processes related to ABA signaling (GO:0009687) and the cellular response to oxygen-containing compounds (GO:1901701) were also enriched.

**Figure 2:**
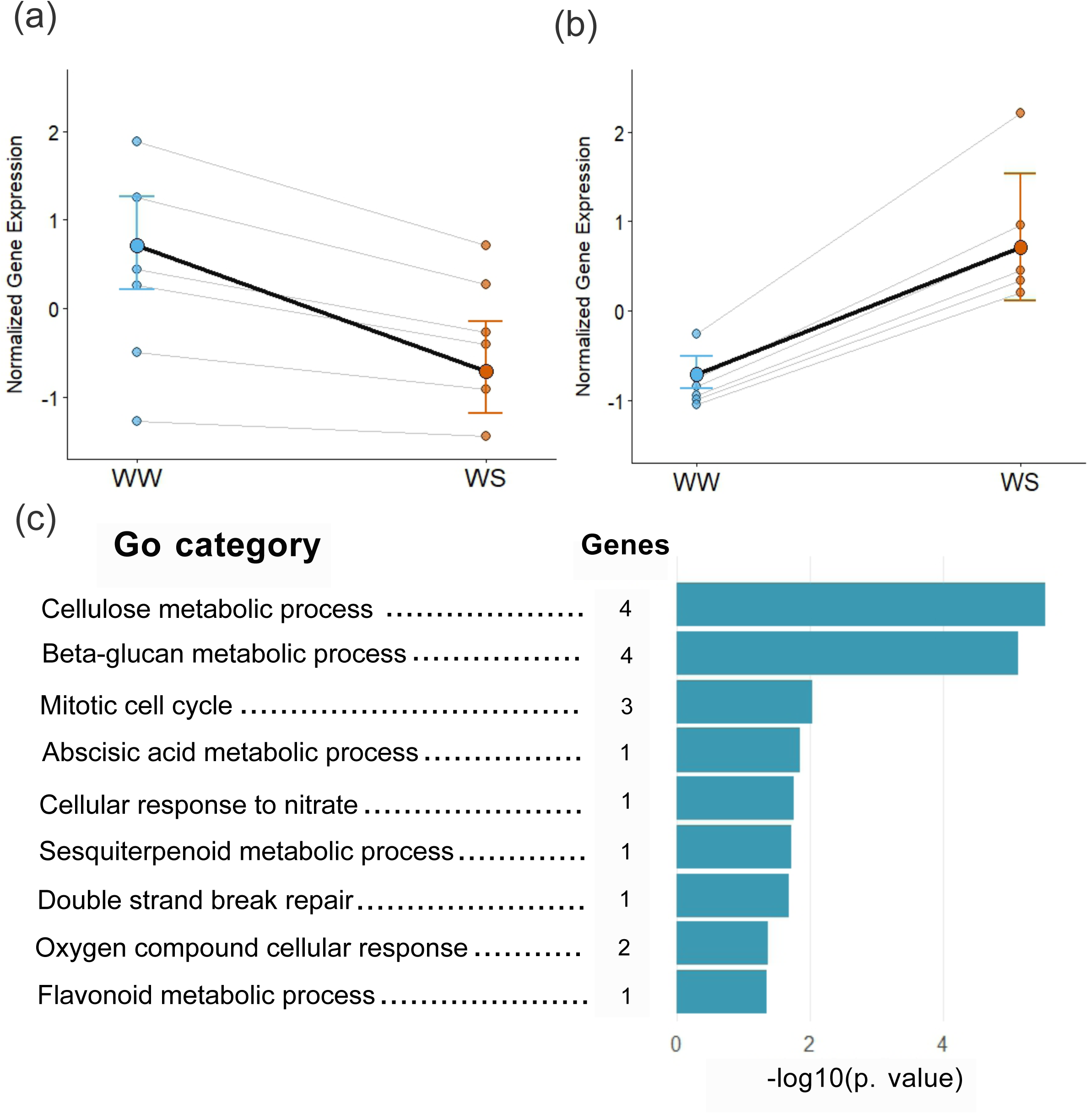
Plastic genes in response to water limitation. (a) Expression of the plastic gene “DIABRO_002784” downregulated under drought. (b) Expression of the plastic gene “DIABRO_023861” upregulated under drought. (c) Gene Ontology (GO) enrichment analysis of the 57 plastic genes, showing the number of genes associated with each GO term. The length of the bars represents the –log10(adjusted p-value) of the enrichment for each term.

### 3.3 Canalization and decanalization in response to drought stress

Regarding plasticity in expression variance, most of the genes showed no significant change in dispersion (LRT *p* ≥ 0.01) and were therefore considered canalized across conditions. Among the genes with significant LRT results, 38 genes showed lower dispersion under drought and were classified as canalized under drought, whereas 182 genes showed higher dispersion under drought and were classified as decanalized under drought (Figure 3 a,b). Functional enrichment analysis (topGO, FDR ≤ 0.05) revealed that decanalized genes under drought were enriched in photosynthesis (GO:0015979) and light reaction processes (GO:0019684), as well as primary amino acid metabolism (GO:0006525). In contrast, canalized genes under drought stress were enriched in immune responses (GO:0002253), regulation of mRNA processing (GO:0050684), and post-transcriptional regulation (GO:0016441).

**Figure 3:**
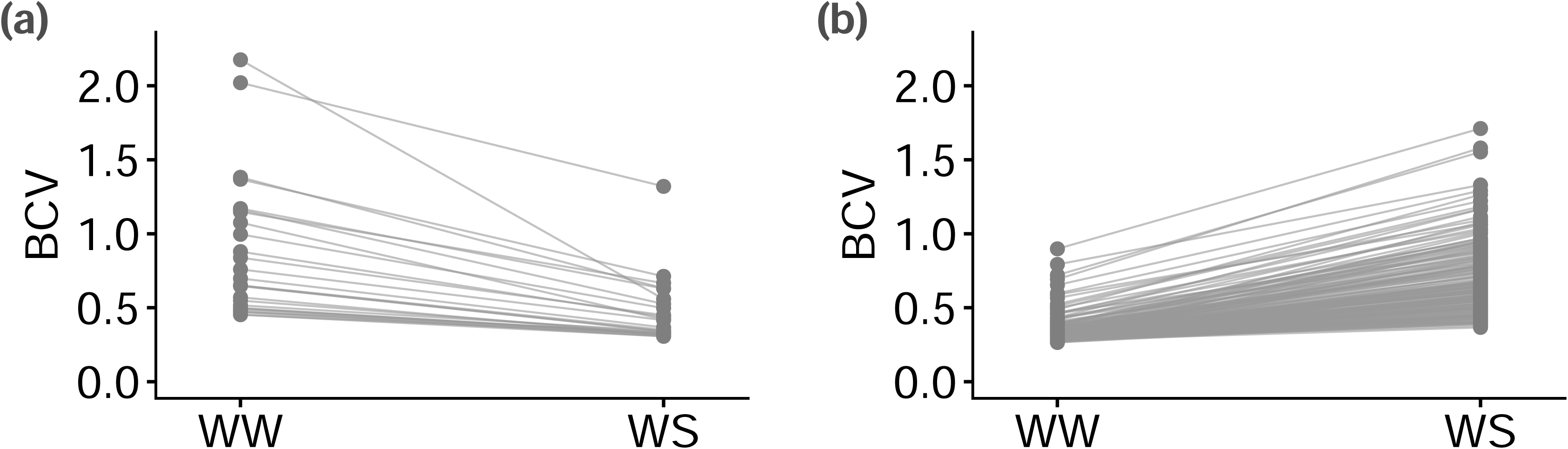
Biological coefficient of variation of (a) underdispersed genes in drought condition and (b) overdispersed genes under drought.

### 3.4 Gene Coexpression and Gene Regulation Networks

The signed-hybrid GCN comprised 31,849 genes, of which 26,274 (82.5%) were assigned to 207 biologically meaningful modules, excluding the unassigned (‘grey’) module that grouped genes with weak or inconsistent correlations. Module sizes ranged from 50 to 1,244 genes. Eigengene–drought correlations identified 15 significant drought-responsive modules (r = 0.58–0.80; p < 0.05; blue, chocolate4, darkgoldenrod1, darkorchid3, darkturquoise, ivory, lavenderblush3, mediumpurple1, midnightblue, navajowhite4, palegreen4, plum3, purple2, sienna, yellow2; Figure 4a). These modules exhibited a limited but clearly defined hub structure, with the number of hub genes ranging from 20 in darkgoldenrod1 to two hub genes in ivory module (Supplementary Table 4). Only two plastic genes belonged to these drought-responsive modules, DIABRO_037551 (unknown protein) in darkgoldenrod1 and DIABRO_023861 (ATCESA7-like) in ivory. Among the plastic genes, four also acted as highly connected hubs in modules not directly associated with drought: DIABRO_012336 (RD22-like, deepskyblue4), DIABRO_029083 (splicing factor 3B subunit 3-like, navajowhite4), DIABRO_042066 (5’ exonuclease Apollo-like, orange3), and DIABRO_013748 (T2 family ribonuclease, skyblue1). Their strongest co-expression links (top 1% adjacency) targeted genes in drought-responsive modules, indicating that these plastic hubs act as inter-modular connectors. For example, DIABRO_012336 (deepskyblue4) connected mainly to modules blue and purple2, whereas DIABRO_013748 (skyblue1) linked to lavenderblush3 and ivory, all significantly associated with drought. Functional enrichment analyses revealed that the 15 drought-associated modules were significantly enriched in four broad functional categories of stress-related responses (Figure 4) including general stress-response pathways, transcriptional and translational regulatory, osmotic adjustment processes, cell-wall modifications and hormone-related processes.

**Figure 4:**
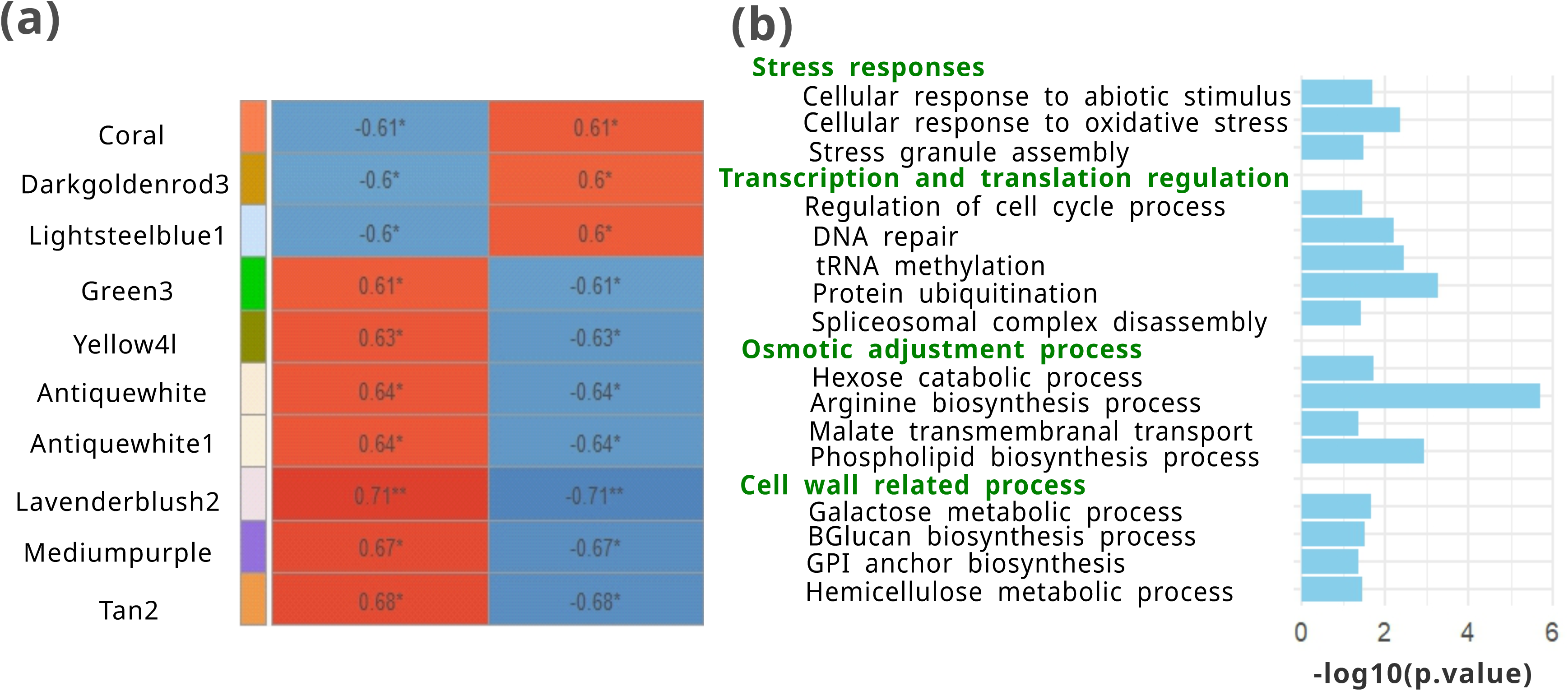
Drought-responsive gene co-expression modules. (a) Heatmap showing the biweight midcorrelation (bicor) coefficients between drought-responsive gene co-expression network modules (rows, identified by color) and water treatment. Asterisks indicate statistically significant correlations (*p* < 0.05) (b) Summary of the enriched gene ontology (GO) categories for the drought-responsive modules. The length of the bars shows the -log (adjusted *p*-value) of the category’s enrichment.

Additionally, the initial GRN comprised 55.1 million predicted regulatory interactions. Applying the topology- and stability-based filtering identified a robust subnetwork of ∼2.43 million high-confidence edges linking 1,362 transcription factors to 30,469 target genes, which was used for all downstream analyses. Analysis of out-degree in the filtered GRN identified several transcription factors with exceptionally broad regulatory reach. The strongest global regulators (Supplementary Table 5) included DIABRO_003068 (AP2/B3), DIABRO_029891 (AGAMOUS65-like), and DIABRO_028208 (ARF16-like). All three TFs had very high out-degree (>2,700 targets) and, as typical in highly directed GRNs, showed zero betweenness and maximal directed closeness, indicating broad direct targeting rather than mediator roles.

Of the identified 57 plastic genes, 55 genes (96%) had at least one predicted transcription factor, whereas only two genes (DIABRO_008569, DIABRO_017591) lacked these regulatory inputs. The plastic gene set received a total of 5,073 TF–gene edges, involving 1,295 distinct TFs, indicating that plasticity is controlled by a highly distributed regulatory architecture. All 55 plastic genes were simultaneously controlled by 15 or more TFs, often combining factors from different TF families. These hyper-connected plastic genes include DIABRO_030906 (CTPS1-like gene), DIABRO_032623 (SAP12-like), DIABRO_043572 (RNA-directed DNA polymerase), DIABRO_005722 (CYP72A15-like), DIABRO_034718 (FLA11-like) and others (Supplementary Table 3), suggesting that coordinated inputs from several stress-responsive regulators contribute to plastic expression changes. A subset of drought-associated TFs showed direct targeting of plastic genes. In total, 42 drought TFs (AREB/ABF, DOF, DREB, GRF, HSF, NAC, WRKY and bZIP families) regulated at least one plastic gene. The dTFs with the highest number of plastic targets were DIABRO_022782 (ATAF1, NAC family; 14 plastic targets), DIABRO_043306 (ERF34, DREB family; 9 targets) and DIABRO_020233 (ERF43, DREB; 8 targets). Additional high-ranking dTFs included DIABRO_027875 (TINY2, DREB), DIABRO_034930 (WRKY33), DIABRO_015516 (AREB3), DIABRO_016491 (GRF7), DIABRO_016693 (ERF58/RAP2.4) and DIABRO_011032 (WRKY2) each regulating 6–8 plastic genes (Supplementary Table 6).

In addition, the integration of GRN with the GCN modules associated with drought stress showed that these modules receive extensive regulatory input, with total incoming edges ranging from 3,763 (darkturquoise) to 18,636 (darkgoldenrod1). When normalized by module size, the regulatory burden was more similar across modules (ranging 70.2–84.3 edges per gene). Weighted load values were also comparable (weight per gene 0.304–0.338), indicating homogeneous TF-driven control despite differences in module size. Enrichment analysis identified numerous TFs significantly associated with each drought-responsive module (p-adjusted ≤ 0.05, ≥2 targets). Across modules, the strongest signals consistently involved TFs with many targets inside the module and moderate global degree (Supplementary Table 7). For example, in the antiquewhite module (553 genes), the top enriched TFs were DIABRO_018813 (WRKY21-like), DIABRO_001330 (WRKY20-like), DIABRO_022736 (trihelix transcription factor), DIABRO_018620 (RGA1-like), and DIABRO_006313 (TFIIIA), each with 99–103 module targets and highly significant enrichment (p-adjusted < 0.001). Equivalent TF rankings were recovered independently using the GRN-based counts of module targets (top 5 per module shown in Supplementary Table 5). Drought TFs previously associated with drought responses were also recovered among the enriched regulators. These 25 dTFs collectively targeted the 15 drought-associated modules, indicating broad but structured regulatory engagement (Figure 5). Their functional annotations revealed that both ABA-dependent and ABA-independent pathways contribute to the control of these modules. Within the ABA-dependent route, AREB/ABF (e.g., AREB3), NAC (e.g., ATAF1) and WRK (e.g. WRK40) showed strong enrichment in several modules, consistent with their role in ABA-mediated activation of stress-responsive genes. In parallel, multiple ABA-independent TF families were also prominent, including DREB, GRF, and HSF, which typically regulate dehydration, ethylene-related signaling, osmotic adjustment, and ROS protection. This combination shows that drought-responsive modules are shaped by TFs acting through both hormonal and hormone-independent routes, with several modules receiving contributions from TFs of both classes.

**Figure 5:**
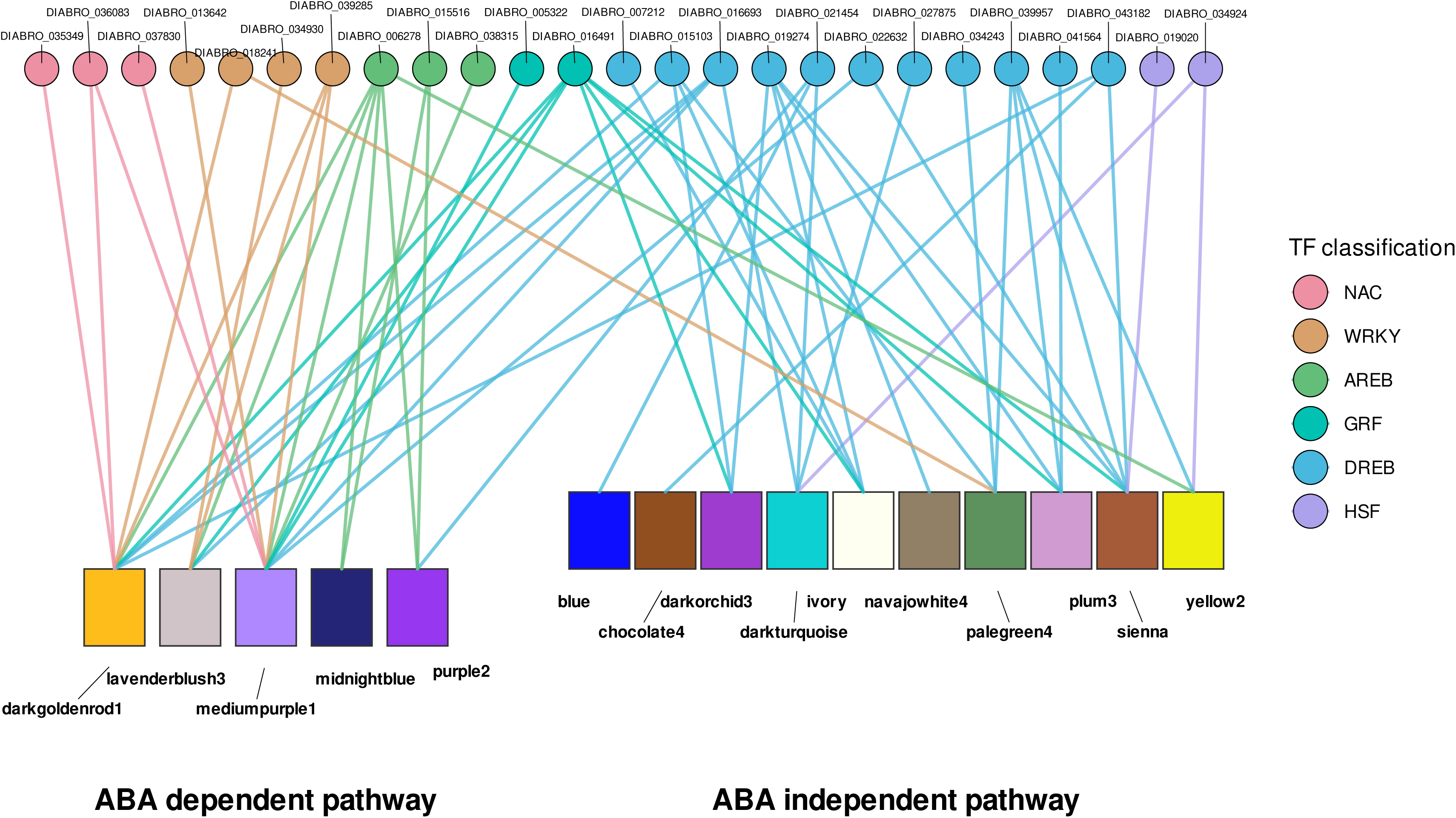
Transcription factor regulating drought-associated modules, including both ABA-dependent and ABA-independent pathways. Transcription factors are colored according to their family (NAC, WRKY, AREB, GRF, DREB, and HSF), while modules are grouped according to their involvement in ABA-dependent (left) and ABA-independent (right) signaling pathways.

## 4. Discussion

### 4.1 Phenotypic plasticity in functional traits

Phenotypic plasticity is expected to play a central role in shaping plant responses to environmental stressors, specifically in the Mediterranean habitats characterized by prolonged periods of extreme drought (Bongers et al., 2017; Ramos-Muñoz et al., 2024). Our results, however, show that *D. inoxianus* exhibited low phenotypic plasticity in response to drought across the eco-physiological traits measured. Despite this overall low plasticity, the most plastic traits were Ψw and Ψπ. Both parameters are key components of plant water relations (Kirkham, 2023) and are among the first to be directly affected by soil water depletion. In the case of Ψπ, its high plasticity indicates that osmotic adjustment (OA) is occurring, a process through which plants accumulate compatible solutes to lower their osmotic potential while maintaining turgor and sustaining physiological functioning under water deficit (Serraj & Sinclair, 2002; Turner, 2018). Enhanced drought resistance through osmotic adjustment has also been reported in many Mediterranean species (Aranda et al., 2004; Martìnez et al., 2004) and even in species of the genus *Dianthus*, such as *D. caryophyllus* (Álvarez et al., 2009; Kwon et al., 2019).

The plastic adjustment in primary water-relation traits appears to buffer the impact of drought on traits more directly linked to plant performance, such as RWC and Fv/Fm, which exhibit low plasticity. The fact that both traits remained largely stable under water stress indicates that the plant can maintain turgor and photosystem II functionality, thereby preserving the performance of the photosynthetic machinery. This suggests that *D. inoxianus* shows a high tolerance to drought stress, consistent with previous studies reporting this drought adaptation based on physiological traits (López-Jurado et al., 2016) and functional traits (López-Jurado et al., 2022), as well as a water-acquisitive ecological strategy in response to drought (López-Jurado et al., 2022). The limited contribution of trait plasticity to drought tolerance may suggest that local adaptation predominantly drives these responses, supporting the maintenance of stable phenotypic traits under water stress (Maghrebi et al., 2025). Previous studies emphasize that, initially, phenotypic plasticity may facilitate the differentiation into unoccupied niches with extreme habitats (Fitzpatrick, 2012). However, once established in a stable environment, selection will reduce plasticity by favoring genotypes that exhibit optimal phenotypes in that given environment, particularly if phenotypic plasticity entails costs (Grenier et al., 2016). In this framework, *D. inoxianus*, which is adapted to dry environments, likely experienced a reduction of its ancestral plasticity during adaptation (López-Jurado et al., 2019). Its habitat is characterized by strong seasonal drought, with long, dry summers and winters with regular rainfall, generating an environment that is variable yet highly predictable. Under such conditions, extreme plasticity is not expected to be favored but rather, selection should promote an intermediate level of plasticity. Consequently, moderate phenotypic plasticity in response to water limitation is the most likely adaptive strategy, allowing individuals to adjust their phenotype in response to environmental cues (Sekajova et al., 2025). This context, in which adaptive responses are shaped by predictable environmental cues, provides a conceptual framework that helps clarify the role of phenotypic plasticity within the genus *Dianthus*. For instance, *D. superbus* subsp. *sylvestris* appears highly plastic in response to variation in altitude and temperature, whereas *D. superbus* subsp*. alpestri* shows ecotypic differentiation rather than plastic responses (Hardion et al., 2020).

### 4.2 Transcriptomic plasticity genes

Consistent with the low phenotypic plasticity observed in its traits, *D. inoxianus* also exhibited limited transcriptional plasticity, with only 57 plastic genes identified. The expression of most genes remains largely unchanged under water deficit. In contrast to many transcriptomic studies reporting extensive genome-wide transcriptional reprogramming under drought, frequently involving thousands of differentially expressed genes (e.g. Ma et al., 2024; Mun et al., 2017), the drought response of *D. inoxianus* appears to involve a limited set of transcriptionally plastic genes. This pattern suggests that drought tolerance in this species relies on specific and targeted transcriptional changes, rather than on broad transcriptomic shifts (Steele et al., 2025).

Despite their relatively small number, the 57 plastic genes identified in our study are enriched in multiple functions associated with drought response. The most significant enrichment function was the modification of cell wall components, a process that enables plants to adjust wall extensibility and thereby maintain optimal turgor under water deficit (Moore et al., 2008). Within this functional category, we found a key plastic gene DIABRO_023861 (ATCESA7-like) which was also found to be part of a module associated with drought. This gene belongs to the Cellulose synthase A family *(CESA)*, associated with cellulose biosynthesis. Previous studies have shown that reduced expression of CESA genes results in weaker secondary walls, leading to xylem collapse under water stress (Chen et al., 2005; Turner & Somerville, 1997). In our case, the plastic up-regulation of this gene under drought likely contributes to thicker and mechanically reinforced cell walls, thereby supporting more efficient water transport and enhancing drought tolerance. Moreover, Clifford (1998) reported that in species which show osmotic adjustment (as occurs in *D. inoxianus*) maintaining more rigid cell walls may be essential to preserve tissue integrity. Therefore, in this context, gene expression plasticity may contribute to enhanced wall integrity and mitigates the detrimental effects of water deficit.

### 4.3 Gene Coexpression and Gene Regulation Networks

Several of the plastic genes identified in this study played a key role within the gene co-expression, highlighting the value of integrating transcriptional plasticity with network-based approaches to better understand drought responses. Specifically, two plastic genes, DIABRO_037551 and DIABRO_023861 (ATCESA7-like), were assigned to drought-associated modules. This limited number suggests that transcriptional plasticity is concentrated in a few key modules rather than being broadly distributed across the network. In addition, beyond plastic genes directly associated with drought-sensitive modules, we found hub plastic genes from other modules which showed strong interactions with drought-associated modules. One illustrative is the DIABRO_029083 gene (splicing factor 3B subunit 3-like). Although there is no previous literature on the role of this specific factor under drought stress, there are a few studies that have highlighted the broader importance of the spliceosome in the responses to water stress (Song et al., 2020; Thatcher et al., 2016). Specifically, the spliceosome triggers numerous isoform changes, thus modifying the function of proteins involved in molecular drought responses. Furthermore, by acting as a hub with strong interactions with drought-associated modules, may function as inter-modular connector linking different drought-responsive pathways.

The gene regulation network suggests that transcriptional control plays a central role in shaping plastic responses, as most plastic genes appear to be under regulatory influence. Among the most strongly regulated plastic genes, we identified DIABRO_030906, a CTPS1-like gene involved in de novo nucleotide synthesis, whose expression was positively regulated under drought conditions. This pattern is consistent with previous findings in *Arabidopsis thaliana*, where CTPS homologues (CTPS1–4) were strongly induced under water stress, and knockout mutants display reduced drought tolerance and poorer recovery after rewatering (Krämer et al., 2022). In our case, the strong and inducible transcriptional regulation of the plastic gene DIABRO_030906, under drought stress suggests an active contribution to drought tolerance and recovery ability. Another plastic gene with a strong regulation was DIABRO_043572, a retrotransposon-derived RNA-directed DNA polymerase that changed its expression under drought and exhibited a high regulatory indegree in the gene regulatory network. This suggests that drought-responsive regulatory programs also impinge on transposable element activity, consistent with stress-induced changes in epigenetic control of TEs (Chen et al., 2025; Lanciano & Mirouze, 2018).

Many plastic genes were regulated by dTF families that are well documented for their involvement in drought stress responses (Manna et al., 2021). Among these, we identified DIABRO_022782 (ATAF1), a transcription factor belonging to the NAC family. ATAF1 is strongly induced by dehydration and abscisic acid (ABA). However, empirical evidence from *Arabidopsis thaliana* indicates that its contribution to stress tolerance is context-dependent. ATAF1-deficient mutants show enhanced recovery after drought (Lu et al., 2007), whereas overexpression of this transcription factor results in increased drought tolerance but is accompanied by pronounced reduced growth (Wu et al., 2009). Another example of dTFs that regulate plastic genes could be WRKY33 and WRKY2. The WRKY family is a large family that includes different TFs that coordinate the response to abiotic and biotic stress through complex regulatory cascades, as studied in *A. thaliana* or *O. sativa* (Banerjee & Roychoudhury, 2015). Specifically, in *A. thaliana*, WRKY33 has been reported to regulate the drought tolerance gene, *CesA8*, which (as mentioned before) encodes a member of the cellulose synthase family required for secondary cell wall biosynthesis (Wang et al., 2013). This is consistent with our results, where we found that its ortholog DIABRO_034930 (WRKY33) regulates DIABRO_036764 (cellulose synthase like D4), a gene also involved in cellulose synthesis, indicating that WRKY33 may modulate cell wall-related processes as a mechanism of adaptation to water stress. Altogether, these results highlight that the regulatory control of plastic expression is not driven by a single regulatory hub but emerges from the combined action of numerous TFs, including several key drought-response regulators that converge onto overlapping subsets of plastic genes. This coordination implies that these dTFs directly modulate transcriptional plasticity and could be important targets of natural selection, contributing to adaptive responses to water limitation in species like *D. inoxianus*.

Finally, we elucidated the regulation of gene co-expression network modules. The drought-responsive modules showed a highly coordinated transcriptional control, with contributions from both ABA-dependent and ABA-independent pathways. This suggests that complementary and redundant mechanisms are operating, allowing plants to respond flexibly to water stress. In fact, several studies highlight that the relationships between ABA-dependent and ABA-independent signaling pathways can be complex, involving significant crosstalk (Du et al., 2018; Roychoudhury et al., 2013; Soma et al., 2021). The observed patterns mixing of modules controlled by ABA dependent and independent regulation pathways match the expected division between early dehydration signalling (DREB/ERF/WRKY/NAC/HSF) and ABA-dependent induction (AREB/bZIP) and explain why multiple modules display mixed TF inputs rather than exclusive pathway-specific control. We identified typical ABA-dependent TFs families such as AREB, NAC and WRKY which are well known for their molecular activities in response to water stress (Soma et al., 2021; Yoshida et al., 2010; Yu et al., 2021). Likewise, ABA independent TFs families were also detected, for example the HSF family. This is consistent with previous observations in other *Dianthus* species, where high expression of HSF was reported under water stress conditions (Li et al., 2019). Together, these findings show that multiple regulatory pathways converge to coordinate gene expression, providing flexibility and drought tolerance to *D. inoxianus*.

### 4.4 Canalization and decanalization in response to drought stress

In this study, water stress significantly increased canalization in some genes while decreasing it in others. Genes that became canalized under drought conditions tended to be associated with immune responses, regulation of mRNA processing, and post-transcriptional regulation, suggesting that these processes are critical and must remain stable even under stressful conditions (Flatt, 2005). In particular, the enrichment of mRNA regulatory functions indicates that this process is essential for plant survival under drought by allowing robust and coordinated gene expression during stress.

Furthermore, we identified a larger set of genes showing decanalization in response to drought treatment. The overrepresentation of decanalized genes involved in photosynthesis and amino acid metabolism, suggests that the expression of these processes becomes more variable under drought as they adjust to changing environmental conditions. Instead of being detrimental, such decanalization confers flexibility to these biological processes. Similar patterns of post-stress decanalization have been reported in other studies (Chen et al., 2015), indicating that decanalization releases previously hidden or “cryptic” genetic variation upon which natural selection can act (Flatt, 2005). In this way, decanalization represents a key mechanism of adaptive evolution, as it converts robustness (of canalized genes) into exploitable genetic diversity, enabling rapid evolutionary responses to environmental change.

## 5. Conclusions

Overall, our results indicate that drought tolerance in *Dianthus inoxianus* is primarily supported by low phenotypic and transcriptomic plasticity, reflecting constitutive adaptations that preserve stability in essential physiological traits and regulatory processes, rather than on broad plastic responses. Only a small set of genes showed plastic expression under drought, and these genes were concentrated in key drought-response pathways and occupied central positions within co-expression network. The drought-associated module regulation involved both ABA-dependent and ABA-independent transcription factors, highlighting a targeted and tightly controlled plastic response. Altogether, these findings suggest that drought tolerance in *D. inoxianus* emerges from the maintenance of stable core functions combined with selective molecular flexibility in critical processes, providing a robust strategy for persistence in arid Mediterranean environments. Our study demonstrates how a small set of plastic genes is structurally and functionally integrated into gene co-expression and regulatory networks, offering novel mechanistic insight into how targeted plasticity contributes to drought tolerance in preadapted Mediterranean species.

## Data availability statement

We have hosted the scripts and data used in this project in a public repository archived at GitFront https://gitfront.io/r/Albarp13/cUvmuP4mroZc/Dianthus-inoxianus-Plasticity/

## Acknowledgements

We thank MF, CM, JPA and AT for contributing to the experiment set-up. As well as, to the University of Seville Greenhouse and Herbarium General Research Services (CITIUS) for assistance and providing facilities and equipment.

## Additional Information

The authors declare no conflict of interest.

## Notes

### Competing Interest Statement

The authors have declared no competing interest.

### Funding

- Ministerio de Ciencia, Tecnología e Innovación: PDI2023-147752NBI00
- FEDER funds, Andalusian Government and University of Seville: FEDER-US-1381232
- Spanish Ministerio de Universidades FPU predoctoral grant: FPU22/00818

