## Supplementary figures and images for "Eco-physiological and transcriptomic plasticity of *Dianthus inoxianus* in response to drought"

### Figure S1

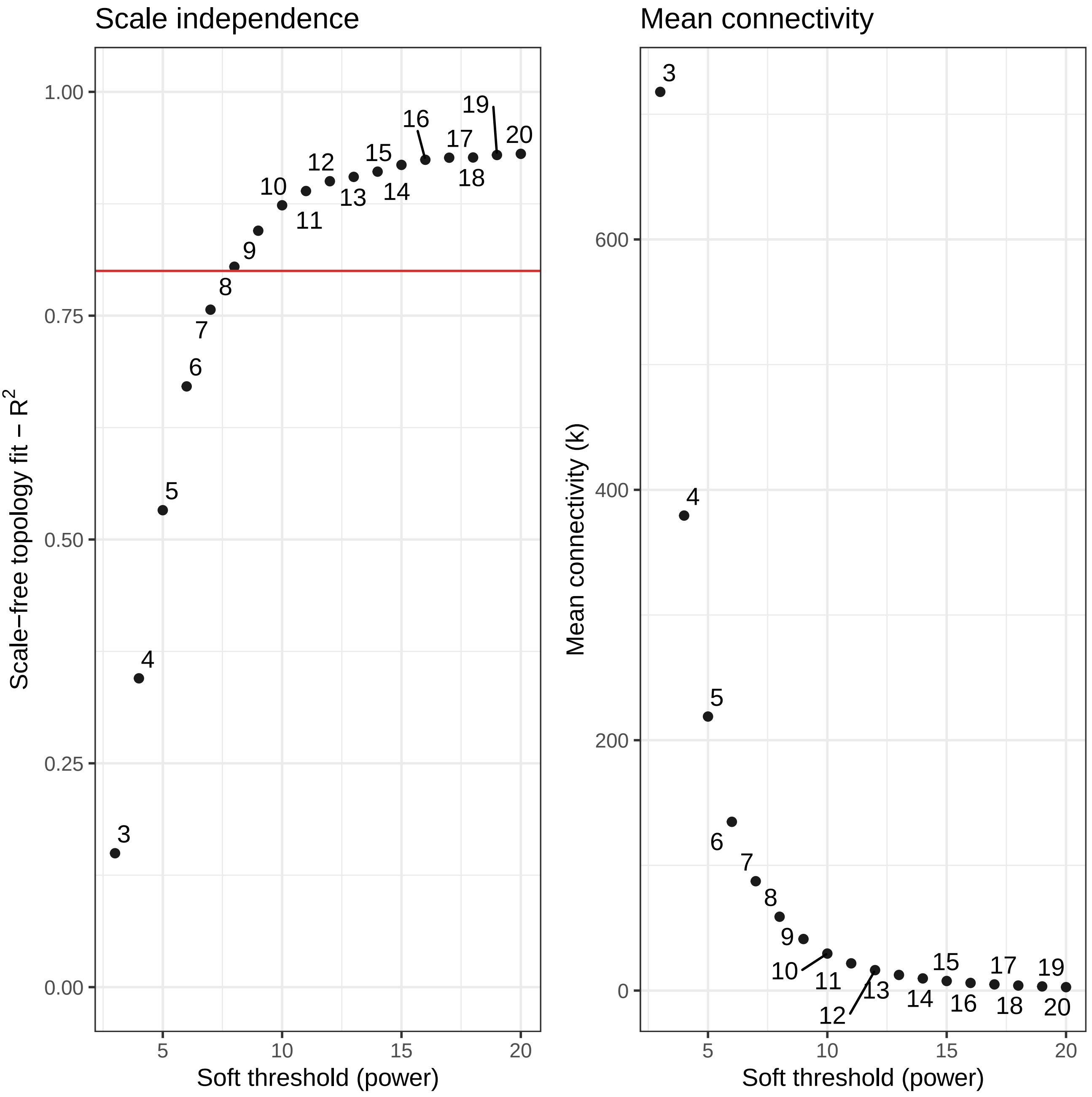
